# Induction of neutralising antibodies against conserved rhinovirus capsid protein VP4 depends on presenting VP4 in a virus-like conformation

**DOI:** 10.1101/2024.06.28.601138

**Authors:** James T. Kelly, Giann K. Dellosa, Joseph Newman, Rory A. Hills, Adel A. M. Mohamed, Julia Aniscenko, Sebastain L. Johnston, Tobias J. Tuthill

## Abstract

Infection with rhinovirus (RV) is associated with significant morbidity and hospitalisation in people with chronic lung disease. At present there is no approved RV vaccine or antiviral. There are approximately 180 RV serotypes, classified into 3 species (RVA, RVB, RVC). There is no cross-protection between serotypes because of high diversity in immunodominant antigenic sites. This makes it impractical to create a broadly protective vaccine using traditional methods, involving whole capsids. A more feasible strategy is to direct the immune response towards conserved but less dominant epitopes. The capsid protein VP4 is highly conserved within each RV genotype and some antibodies that target VP4 are neutralising. This makes VP4 vulnerable to antibodies and a promising vaccine target.

Here we investigate the ability of RV VP4 N-terminal peptides presented on different display systems to initiate a neutralising immune response in mice. We compared three different-sized VP4 peptides (spanning residues, 1-15, 1-30 and 1-45) displayed on two different display systems-SpyCatcher Virus-like particles (VLPs) or Keyhole Limpet Hemocyanin (KLH). Overlapping regions of VP4 were antigenically different when presented on different platforms and in peptides with different length. The conformation of the 1-15 region of VP4 was critical for inducing antibodies that were both neutralising and able to bind VP4 in the context of the virus particle. Therefore, correctly displayed VP4 peptides can recapitulate virus-like antigenicity. These findings improve our understanding of VP4 antigenicity and will inform the design of future RV vaccines. This work could also impact the design of other peptide vaccines, since a variable antigenic conformation is a common characteristic of pathogen-derived peptide targets.

**Impact statement:** Rhinovirus (RV) is a highly prevalent respiratory virus that is a frequent cause of common cold symptoms. For people with chronic lung diseases such as asthma, infection can cause significant worsening of disease, leading to increased suffering and hospitalisation. There are currently no vaccines or anti-viral drugs that target RV infections. Development of a RV vaccine would be hugely beneficial to suffers of chronic lung diseases.

Attempts to develop a broadly protective RV vaccine have been hindered by the presence of over 180 serotypes which have little or no natural cross-reactivity. Previous studies have identified the highly conserved VP4 epitope as a potential target for broadly protective vaccines. However, there has been little progress in evaluating its effectiveness as a vaccine candidate.

In this study we assessed the ability of different RV VP4 immunogens to induce neutralising antibodies in mice. We demonstrated that antigenic conformation of VP4 varies greatly between immunogens and presentation of the first 15 N-terminal residues in a virus-like conformation is important for generating neutralising antibodies.

This knowledge could help to maximise the production of neutralising VP4-specific antibodies in future studies which could ultimately lead to a universal RV vaccine.

## Introduction

Rhinovirus (RV) predominantly causes a mild self-limiting illness in healthy individuals. However, infection is associated with significant morbidity in those with chronic lung diseases such as asthma (Johnston et al., 1995) and chronic obstructive pulmonary disease (COPD) exacerbations (Seemungal et al., 2001). RV was present in 42% of hospitalisations due to respiratory tract infections (Aponte et al., 2015). Therefore, treatment and reducing the spread of RV would be hugely beneficial to sufferers of lung disease. At present there is no approved RV vaccine or antiviral.

RVs are members of the enterovirus genus in the picornavirus family. There are 3 RV species known as RVA, RVB and RVC. RV species are further distinguished by receptor usage. RVA uses either Intercellular Adhesion Molecule 1 (ICAM-1) or low density lipoprotein receptor (LDLR). RVB uses ICAM-1. RVC uses cadherin-related family member 3 (CDHR3) (Bochkov & Gern, 2016). Amongst these species there are ∼180 serotypes, which have little or no natural cross-reactivity (McLean, 2020). Attempts to generate cross-reactive protective/neutralising responses using inactivated virus vaccines have only been successful by immunisation with a multivalent vaccine, with one such study successfully creating a 50-valent response in rhesus macaques (Lee et al., 2016). However, these vaccines do not target a conserved epitope, so that the observed neutralisation in multivalent RV vaccines is predominantly limited to the serotypes found in the vaccine formulation (Cooney et al., 1973; Hamory et al., 1975; Lee et al., 2016). This means that an effective multivalent RV vaccine may need to include all 180 serotypes, which would not be feasible.

There is low cross-protection between virus serotypes because the major Neutralising Immunogenic (NIm) sites are formed by discontinuous epitopes on the capsid surface that are highly variable between serotypes (Narean et al., 2019). A more feasible strategy to create a broadly protective RV vaccine would therefore use a vaccine formulation that directs the immune response towards conserved epitopes. Highly conserved regions within RV and other picornaviruses have been identified as binding sites for neutralising antibodies, such as the internal capsid protein VP4. Immunisation against full length VP4 or peptides from VP4 has generated neutralising antibodies against several different picornaviruses including RV, Poliovirus and Enterovirus A71 (EV-A71) (Katpally et al., 2009; Li et al., 1994; Panjwani et al., 2016; Phanthong et al., 2020; Zhao et al., 2013). RV VP4 antibodies neutralised multiple diverse serotypes: antibodies raised against VP4 of RV-B14 neutralised RV-B14 as well as RV-A16 and RV-A29 (Katpally et al., 2009).

The mechanism of neutralisation for these VP4 antibodies is likely through the prevention of interaction between VP4 and the endosomal membrane (Panjwani et al., 2016). Under physiological conditions, VP4 is transiently exposed at the surface of the virus, this process is known as capsid breathing (Katpally et al., 2009; Lewis et al., 1998). Picornavirus VP4 plays an essential role during virus entry: when in contact with a membrane such as the endosome, VP4 permeabilises this membrane to form a size-selective pore (Danthi et al., 2003; Davis et al., 2008; Kelly et al., 2022; Panjwani et al., 2014; Shukla et al., 2014). Formation of this pore is expected to facilitate the viral genome’s exit from the capsid across the endosomal membrane and into the cytoplasm, where the genome can then replicate (Panjwani et al., 2014). Antibodies raised against the first 16 N-terminal residues of VP4 from RV-A16 can block viral pore formation in model membranes and were capable of neutralising infection (Panjwani et al., 2016). Further evidence of the ability of VP4 to induce protective responses can found from *in vivo* studies. Inoculation of suckling mice with sera from mice immunised with a Hepatitis B Virus core (HepB core) fusion protein containing the N-terminal 20 residues of VP4 from EV-A71 protected from EV-A71 infection (Zhao et al., 2013).

These studies show that VP4 has the potential to act as a broadly protective vaccine for RV. However, not all antibodies that target VP4 are neutralising. When antibodies were raised against peptides corresponding to the N-terminal 16 or C-terminal 16 amino acids of RV-A16 VP4, neutralisation was only observed for antibodies raised against the N-terminal peptide (Panjwani et al., 2016). Antibodies raised against peptides corresponding to the N-terminal 24 and 30 amino acids of RV-B14 VP4 and the N-terminal 20 amino acids of EV-A71 VP4 are also neutralising (Katpally et al., 2009; Zhao et al., 2013). These results suggest that only activity against the N-terminus of VP4 is required for neutralisation.

There is also evidence that the length of VP4 peptide influences the antigenic conformation, which could influence its ability to induce neutralising antibodies (Katpally et al., 2009). Therefore, the length of VP4 used in immunisations must be carefully optimised to ensure that the most neutralising site is targeted to achieve maximum protection.

In addition to the length of peptide, the method of presentation must also be investigated. Free peptides initiate low levels of immunogenicity (Purcell et al., 2007), whereas displaying antigens on a VLP or carrier protein substantially enhances immunogenicity (Gomes et al., 2017; Hills & Howarth, 2022). There are a variety of different presentation systems available, including the SpyTag/SpyCatcher system(Brune & Howarth, 2018), which was recently shown to generate good immune responses to several different immunogens including the SARS-CoV-2 receptor binding domain (Tan et al., 2021). In this system the VLP subunit is genetically fused to the SpyCatcher003 domain, which allows the VLP to be decorated with SpyTag-fused antigens through spontaneous formation of covalent isopeptide bonds (Tan et al., 2021). The well-established Keyhole Limpet Hemocyanin (KLH) and HepB core assemblies for antigen presentation have already been demonstrated to induce immune responses against picornavirus VP4 peptides (Katpally et al., 2009; Panjwani et al., 2016; Swanson et al., 2021; Zhao et al., 2013).

In the present study we compared the ability of KLH and SpyTag/SpyCatcher presentation systems to display RV-A16 N-terminal VP4 peptides corresponding to amino acids 1-15, 1-30 and 1-45 and induce an immune response in mice. We mapped which epitopes are bound by the sera bind and assessed the ability to neutralise RV infections. Here we show that both display systems can generate an immune response to various VP4 peptides. We also determine how peptide length and display system influence which part of VP4 is targeted by the antibody response.

## Results

### Different lengths of VP4 peptides can be effectively displayed on mi3 VLP via SpyTag/SpyCatcher

To create a VP4-SpyVLP vaccine candidate, VP4 peptides of different lengths were solid-phase synthesised with a run of 6 lysines as the spacer and then a C-terminal SpyTag (AHIVMVDAYKPTK) (Table 1). The lysines improve the solubility, since VP4 is very hydrophobic. VP4-SpyTag peptides were conjugated to the SpyCatcher003-mi3 VLP (SpyVLP) to generate the VP4-SpyTag:SpyCatcher003-mi3 (VP4-SpyVLP) immunogen (Bruun et al., 2018; Tan et al., 2021). mi3 is a dodecahedral 60-mer nanocage, engineered from the *Thermotoga maritima* aldolase(Bruun et al., 2018; Hsia et al., 2016). Six different peptides were tested, including myristoylated and non-myristoylated VP4 peptides, spanning amino acids 1-15 (1-15-VP4), 1-30 (1-30-VP4) and 1-45 (1-45-VP4) (Table 1).

**Table 1.** Summary of neutralisation at dilution 1 in 2 and 1 in 4.

| Immunisation | Number that are neutralising at 1:2 | Number that are neutralising at 1:4 |
| --- | --- | --- |
| 1-15-VP4-SpyVLP-0.5 | 1/5 | 1/5 |
| 1-30-VP4-SpyVLP-0.5 | 2/5 | 0/5 |
| 1-45-VP4-SpyVLP-0.5 | 1/5 | 0/5 |
| 1-15-VP4-SpyVLP-5 | 5/5 | 5/5 |
| 1-30-VP4-SpyVLP-5 | 1/5 | 0/5 |
| 1-45-VP4-SpyVLP-5 | 0/5 | 0/5 |
| 1-15-VP4-KLH | 3/5 | 0/5 |
| 1-30-VP4-KLH | 2/5 | 0/5 |
| 1-45-VP4-KLH | 1/5 | 0/5 |

Analysis by SDS-PAGE and Coomassie protein staining revealed that non-myristoylated peptides 1-15-VP4 and 1-30-VP4 were conjugated to SpyVLP with high efficiency, as demonstrated by the presence of a single band that is larger than the SpyVLP only control (Figure 1A). For the 1-45-VP4 non-myristoylated peptide, after conjugation there were two prominent bands, indicating lower efficiency of conjugation (Figure 1A). All myristoylated peptides had a low conjugation efficiency as demonstrated by the presence of two prominent bands (Figure 1A).

**Figure 1.**
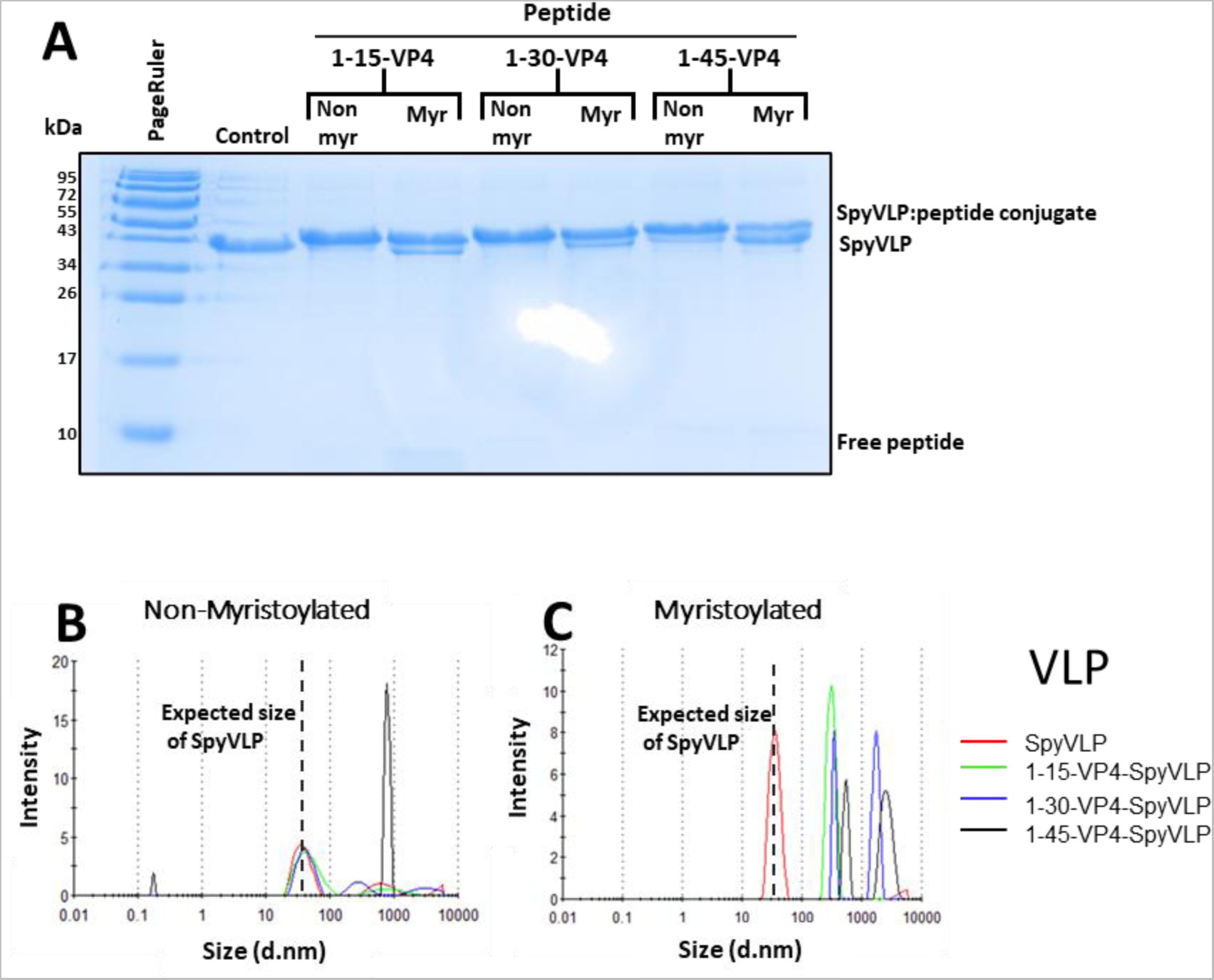
Non-myristoylated SpyTag-VP4 peptides 1-15 and 1-30 are efficiently conjugated to SpyCatcher003-mi3 VLP without aggregation. (A) Conjugation of SpyCatcher003-mi3 with different SpyTag-VP4 peptides, including non-myristoylated (Non myr) and myristoylated (Myr) VP4 peptides, spanning amino acids 1-15, 1-30 and 1-45 from the N-terminus. Reactions were performed at 22 °C overnight and analysed using SDS–PAGE with Coomassie staining. Similar reactions have been repeated multiple times during subsequent preparation of material for immunisations. (B-C) Dynamic light scattering (DLS) characterisation of SpyVLP alone and conjugates 1-15-VP4-SpyVLP, 1-30-VP4-SpyVLP and 1-45-VP4-SpyVLP with or without myristoylation

Analysis by dynamic light scattering (DLS) revealed that non-myristoylated peptides 1-15-VP4 and 1-30-VP4 showed no signs of aggregation following conjugation to SpyVLP. As shown by a peak of the hydrodynamic diameter at 40.9 ± 19.4 nm for 1-15-VP4-SpyVLP and 34.02 ± 15.24 nm for 1-30-VP4-SpyVLP in DLS, as compared to SpyVLP at 35.6 ± 9.5 nm. Due to the small size of the peptides, conjugation does not result in a detectable increase in the hydrodynamic size of the VLP by DLS (Figure 1B). However, conjugation of SpyCatcher003-mi3 VLP to non-myristoylated 1-45-VP4 or the myristoylated peptides 1-15-VP4, 1-30-VP4 and 1-45-VP4 did increase the size of the peak to over 100 nm. This indicates that conjugation between these peptides and the SpyCatcher003-mi3 VLP induces aggregation (Figure 1 B,C). This aggregation could be the cause of the lower conjugation efficiency observed for the myristoylated peptides and the non-myristoylated 1-45-VP4 peptide. The myristoylated VLP conjugates were also observed to precipitate after conjugation. In light of the higher conjugation efficiency, higher solubility and lower aggregation, conjugates of the non-myristoylated peptides were taken forward as test immunogens.

### Sera from mice immunised with 1-15-VP4 peptides conjugated to SpyVLP neutralises RV-A16

In addition to the novel VP4-SpyVLP peptide conjugates described above, myristoylated VP4 peptides were also displayed using the well-established KLH peptide presentation system, as used for previous RV VP4 immunisations in mice and sheep (Katpally et al., 2009; Panjwani et al., 2016).

Mice were immunised with VP4-SpyVLP or VP4-KLH peptide conjugates, presenting VP4 amino acids 1-15, 1-30 and 1-45 from the N-terminus, as follows. Mice (n = 5) were immunised subcutaneously (SC) with either 1-15-VP4-SpyVLP (0.5 or 5 µg), 1-30 VP4-SpyVLP (0.5 or 5 µg), 1-45 VP4-SpyVLP (0.5 or 5 µg), SpyVLP (0.5 or 5 µg), 1-15 VP4-KLH (50 µg), 1-30 VP4-KLH (50 µg), 1-45 VP4-KLH (50 µg), KLH (50 µg). All immunogens were adjuvanted with Magic^TM^ Mouse Adjuvant, which contains immune-stimulatory CpG DNA oligodeoxynucleotides. Mice were boosted with the same dose of immunogen 14 days and 28 days later. Tail bleeds were collected immediately prior to immunisation, 10 days post immunisation and 10 days post boost 1. Terminal bleeds were taken 14 days post boost 2 (Figure 2A). Each bleed was subsequently processed to separate the blood cells from the sera. Sera were then analysed by Virus Neutralisation Test (VNT) to assess the neutralising response from each immunogen. RV-A16 was incubated in the presence of endpoint sera at dilutions from 1 in 2 to 1 in 64, before infecting HeLa-H1 cells. Sera from a sheep immunised with 1-16-VP4-KLH, which has previously been demonstrated to neutralise RV-A16 infections, was included as a positive control (Panjwani et al., 2016). A no sera sample was also included as a negative control. Three days post-infection, cytopathic effect (CPE) was measured by crystal violet staining as a quantitative measure of cell viability (Glanville et al., 2016). Cells incubated in the presence of virus with no sera achieved complete cell death, giving an OD of 0.22. Cells incubated in the presence of 1-16-VP4-KLH positive control sera showed no signs of CPE and had an OD of 1.95 at a 1 in 2 or 1 in 4 dilution; this OD went below OD 0.7 at 1 in 8 and below 0.2 at 1 in 64 (Figure 2B), indicating that the serum is fully neutralising at a 1 in 4 dilution and partially neutralising at a 1 in 8 dilution, which is consistent with previous studies (Panjwani et al., 2016).

**Figure 2.**
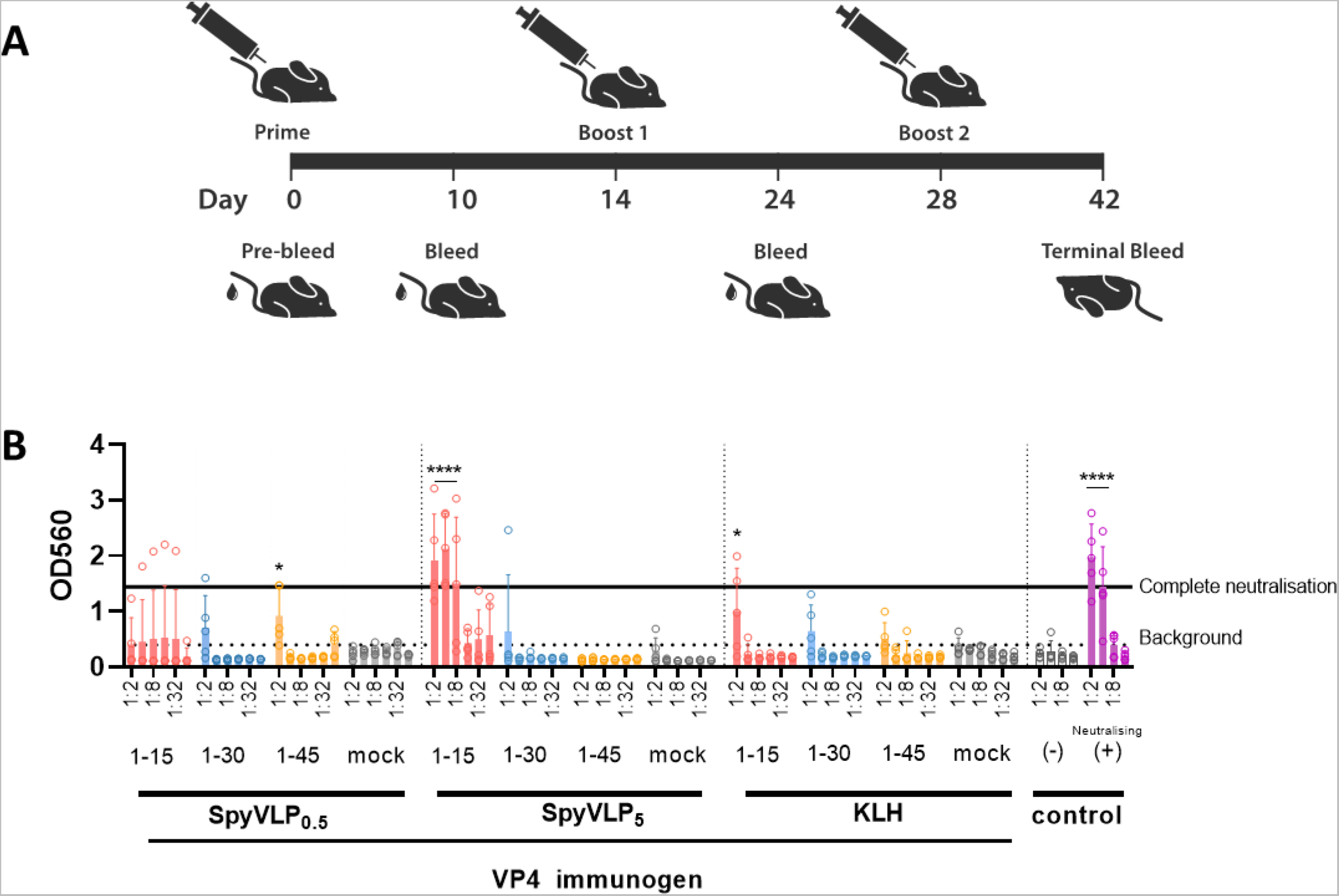
Assessment of neutralising capacity of terminal bleed immune sera using VNT. (A) Schematic diagram of mouse immunisation schedule. (B) VNT was used to assess ability of immune sera to neutralize RV-A16 infections in HeLa H1 cells. Serum dilutions spanned from 1:2 to 1:64. Mouse immune sera samples were 1-15-VP4-SpyVLP-0.5, 1-30 VP4-SpyVLP-0.5, 1-45 VP4-SpyVLP-0.5, SpyVLP-0.5, 1-15-VP4-SpyVLP-5, 1-30 VP4-SpyVLP-5, 1-45 VP4-SpyVLP-5, SpyVLP-5, 1-15 VP4-KLH, 1-30 VP4-KLH, 1-45 VP4-KLH, KLH. Controls included a negative control (PBS) and a neutralising positive control (1-16-VP4-KLH sheep sera from Panjwani et al 2016). CPE was quantified by crystal violet staining and read at OD_560_. Each point corresponds to a single vaccinated mouse from n=5 per vaccination group. Data are shown as mean ± 1 SD. Complete neutralisation is represented by solid blackline, background absorbance is represented by a black dashed line. Statistical analysis was performed using two-way ANOVA and post-hoc analysis was performed using Sidak’s pairwise comparison with a significance threshold of a = 0.05. Comparisons were between corresponding dilution of sera vs mock-vaccinated controls. *p<0.05, ***p<0.005, ****p<0.0005.

Analysis of the endpoint immune sera revealed that sera derived from the mock immunisations with immunogens lacking VP4 (SpyVLP-0.5, SpyVLP-5 or KLH) had a mean OD of 0.27 to 0.35. This is close to the no sera control reading of OD 0.22, which represents full cell death. This indicates that mock sera do not protect cells from infection with RV-A16. However, a single sample in SpyVLP-5 and KLH that had OD of 0.66 and 0.64 respectively at the 1 in 2 dilution, which indicates partial neutralisation (Figure 2B). These sera appear to offer some non-specific background level of neutralisation. Due to the presence of this background neutralisation, we have classified readings above 0.7 OD as neutralising.

For the VP4-containing immunogens, only the 1-15-VP4-SpyVLP-5 gave a neutralising response in all the mice in that group (Figure 2B). In the 1-15-VP4-SpyVLP-0.5 group only 1/5 sera were neutralising, indicating that the 5 µg dose is required to generate a more consistent neutralising response. All 1-15-VP4-SpyVLP sera were effective at neutralisation down to dilutions of 1 in 8.

The 1-15-VP4-KLH group possessed the second highest number of neutralising sera with 3/5. These sera were only capable of neutralising at 1 in 2 dilution. In the remaining groups, the ability to neutralise or partially neutralise was as follows, 2/5 1-30-VP4-SpyVLP-0.5, 1/5 1-30-VP4-SpyVLP-5, 2/5 1-30-VP4-KLH, 1/5 1-45-VP4-SpyVLP-0.5, 0/5 1-45-VP4-SpyVLP-0.5 and 1/5 1-45-VP4-KLH, as summarised in Table 1. No sera from these groups were neutralising below a 1 in 2 dilution.

Of the immunogens tested, 1-15-VP4-SpyVLP-5 produced the strongest and most reliable neutralising response. This level of neutralisation is likely too low to result in a protective response, however a better understanding of this immunogen’s antigenicity could provide information that informs further studies seeking to increase VP4-mediated neutralising response.

### Neutralising sera from mice immunised with 1-15-VP4 immunogens do not react with 1-45-VP4 peptides

Now that we had established that certain end-point sera were neutralising, we wanted to determine how the overall strength of antigen-specific immune response developed during the immunisation experiment. We analysed sera from each bleed time point by ELISA using 1-45-VP4 peptide as the capture antigen, since all immunogens in this study contain a sequence that corresponds with this peptide. Therefore, 1-45-VP4 has the potential to bind all antibodies produced in response to immunogens used in this study.

We found that sera from mice immunised with either SpyVLP-0.5, SpyVLP-5, KLH, 1-15-VP4-SpyVLP-0.5, 1-15-VP4-SpyVLP-5 or 1-15-VP4-KLH did not develop detectable reactivity with the 1-45-VP4 peptide (Figure 3 A,B,C,D,E,F). This is despite the observation that several sera from groups 1-15-VP4-SpyVLP-0.5, 1-15-VP4-SpyVLP-5 or 1-15-VP4-KLH neutralised RV-A16 in VNT (Figure 2B), indicating that they bind part of VP4 that is not accessible in the 1-45-VP4 peptide. This indicates the 1-45 peptide may a adopt a different conformation to the 1-15 peptide.

**Figure 3.**
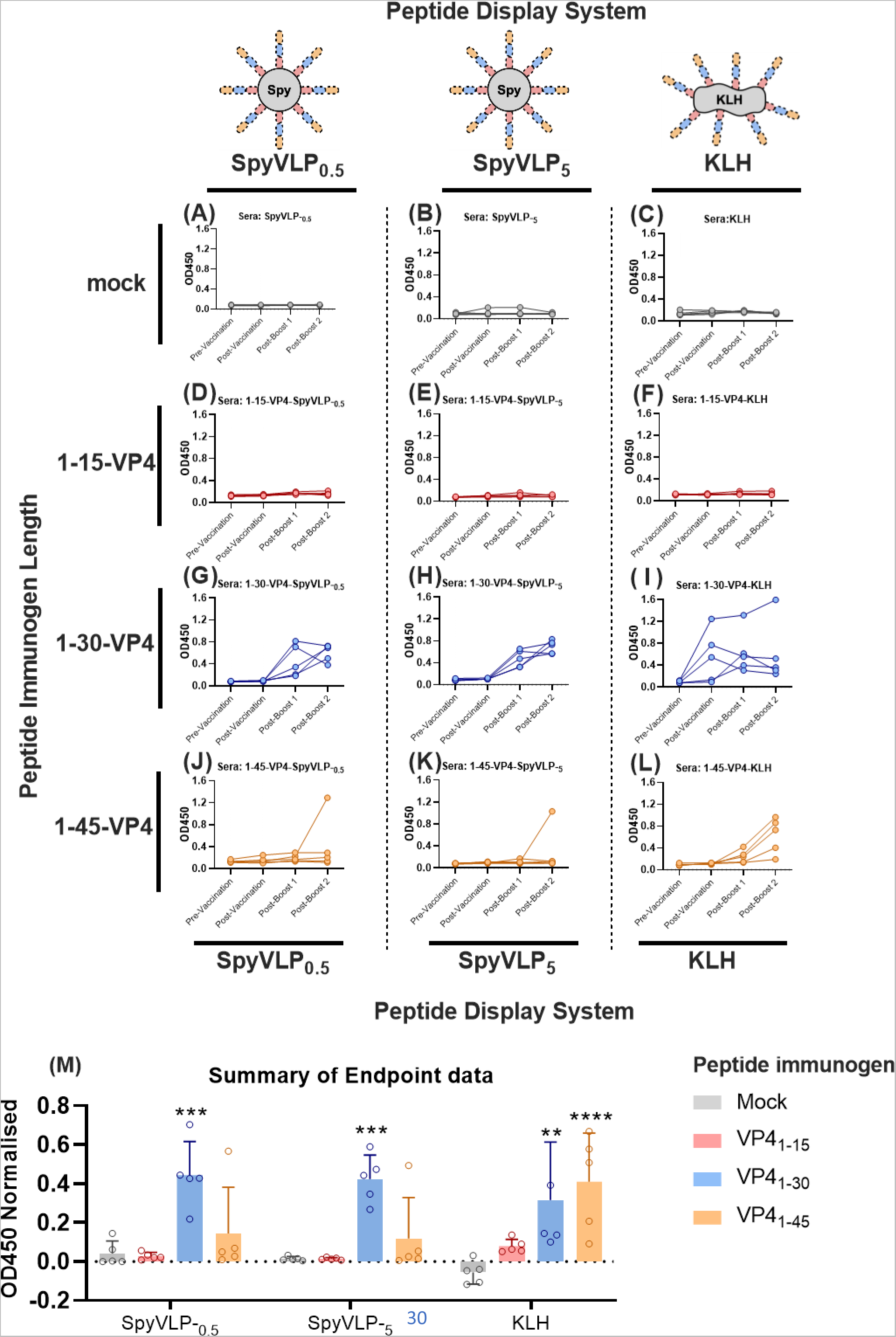
Time-course of reactivity of sera with VP4-1-45 peptide. ELISAs using the VP4-1-45 peptide as a capture were used to assess the reactvity of immune sera. (A-L) Sera corresponding to pre-vaccination, post vaccination, post-boost 1 and post-boost 2 were used at a dilution of 1:160 in ELISA with VP4-1-45 peptide as the capture antigen. Sera from each group was used, SpyVLP-0.5 (A), SpyVLP-5 (B), KLH (C), 1-15-VP4-SpyVLP-0.5 (D), 1-15-VP4-SpyVLP-5 (E), 1-15 VP4-KLH (F), 1-30-VP4-SpyVLP-0.5 (D), 1-30-VP4-SpyVLP-5 (H), 1-30-VP4-KLH (I), 1-45-VP4-SpyVLP-0.5 (J), 1-45-SpyVLP-5 (K), 1-45 VP4-KLH (L). Absorbance was read at OD_450_. All ELISAs were carried out with an anti-mouse IgG secondary antibody. Each line represents an individual mouse. (M) Summary of endpoint (post boost 2) ELISA data in panels A-L. Data were normalised to positive and negative controls. Statistical analysis was performed using two-way ANOVA and post-hoc analysis was performed using Sidak’s pairwise comparison with a significance threshold of a = 0.05. Comparisons were between corresponding dilution of sera vs mock-vaccinated controls. *p<0.05, ***p<0.005, ****p<0.0005.

Analysis of sera from 1-30-VP4-SpyVLP immunised mice, revealed sera from 5/5 mice immunised with 1-30-VP4-SpyVLP-5 were reactive with VP4-1-45 peptide after the 1^st^ boost (Figure 3H). For 3/5 mice, a single boost was sufficient to reach maximum reactivity. For 2/5 mice, reactivity continued to increase after the 2^nd^ boost (Figure 3H). Similar results were observed for mice immunised with 1-30-VP4-SpyVLP-0.5 (Figure 3G). 5/5 mice from this group were reactive with 1-45-VP4 peptide after the 1^st^ boost, while for 2/5 mice the reactivity continued to increase after the 2^nd^ boost (Figure 3G). This indicates that the 5 µg dose of 1-30-VP4-SpyVLP may initiate a slightly superior immune response after the 1^st^ boost.

After immunising with 1-30-VP4-KLH, we found that 5/5 mice had reactivity with the 1-45-VP4 peptide after the 1^st^ boost (Figure 3F). Sera from 1/5 mice had reactivity after the initial immunisation and it had much higher reactivity than all other mice. 4/5 had lower levels of reactivity than the mice immunised with 1-30-VP4-SpyVLP even after the 2^nd^ boost (Figure 3I). This indicates that presentation of the 1-30-VP4 peptide on the SpyVLP produces a superior immune response to the 1-30-VP4 peptide presented on KLH.

Sera from 1/5 mice immunised with 1-45-VP4-SpyVLP-0.5 and 1/5 mice immunised with 1-45-VP4-SpyVLP-5 were reactive with VP4 1-45 peptide; this reactivity only occurred after the 2^nd^ boost (Figure 3 J,K).

Sera from 3/5 mice vaccinated with 1-45-VP4-KLH were reactive with the VP4-1-45 peptide after the 1^st^ boost. Reactivity increased after the 2^nd^ boost, where 4/5 mice sera were reactive (Figure 3L). This demonstrated that presentation of the VP4-1-45 peptide on KLH produces a superior immune response to presentation of VP4-1-45 peptide on the SpyVLP. Although 1-45-VP4-SpyVLP did generate an immune response, it was not consistent or reliable; this may relate to the high levels of aggregation observed in DLS (Figure 1B).

Given that sera from mice immunised with 1-15-VP4 immunogens (3/5 1-15-VP4-KLH, 1/5 1-15-VP4-SpyVLP-0.5 and 5/5 1-15-VP4-SpyVLP-5) neutralised the virus in a VNT, it is surprising that these neutralising 1-15-VP4 immunogens had no detectable reactivity with the 1-45 peptide, even though they produced the strongest levels of neutralisation, 7/15 induced full neutralisation and 2/15 induced partial neutralisation. All 15 immunisations with 1-30-VP4 immunogens produced sera that strongly reacted with 1-45 peptide, but only 1/15 induced full neutralization and only 5/15 induced partial neutralisation. For the 1-45-VP4 immunogens, 7/15 were reacted with the 1-45 peptide and 2/15 were partially neutralising. Therefore, neutralisation and reactivity with the 1-45 peptide do not corelate. This indicates the neutralising epitope may be poorly displayed in 1-30 and 1-45 peptides when presented on an ELISA plate.

### Peptide length and antigen display system influence the site of binding of VP4-specific antibodies

Since the neutralising sera raised against 1-15-VP4-KLH, 1-15-VP4-SpyVLP-0.5 or 1-15-VP4-SpyVLP-5 did not bind the 1-45 peptide, we hypothesized that the longer peptide may adopt a different conformation that influences binding of VP4 antibodies. To determine if the length of the VP4 peptide influences binding, we sought to identify the location of the epitopes on VP4 that each serum reacted with. This was carried out by two different methods using ELISA. Firstly, we assessed the ability of sera to react with equimolar concentrations of all three peptide lengths used in the study (1-15-VP4, 1-30-VP4, 1-45-VP4) using a dilution range of 1 in 80 to 1 in 640. In addition, we mapped the position(s) on VP4 that the sera bound, using a scanning library of 15-mer peptides with 5 amino acid overlaps based on the first 55 N-terminal residues of RV-A16 VP4 and a sera dilution of 1 in 160 (Table ?). Together these experiments allow us to evaluate how peptide length influences antigenicity.

For the mock immunisation groups SpyVLP-0.5, SpyVLP-5 and KLH, no sera from immunisations with these immunogens reacted with any length of VP4 peptide (Figure 4 A,B,C).

**Figure 4.**
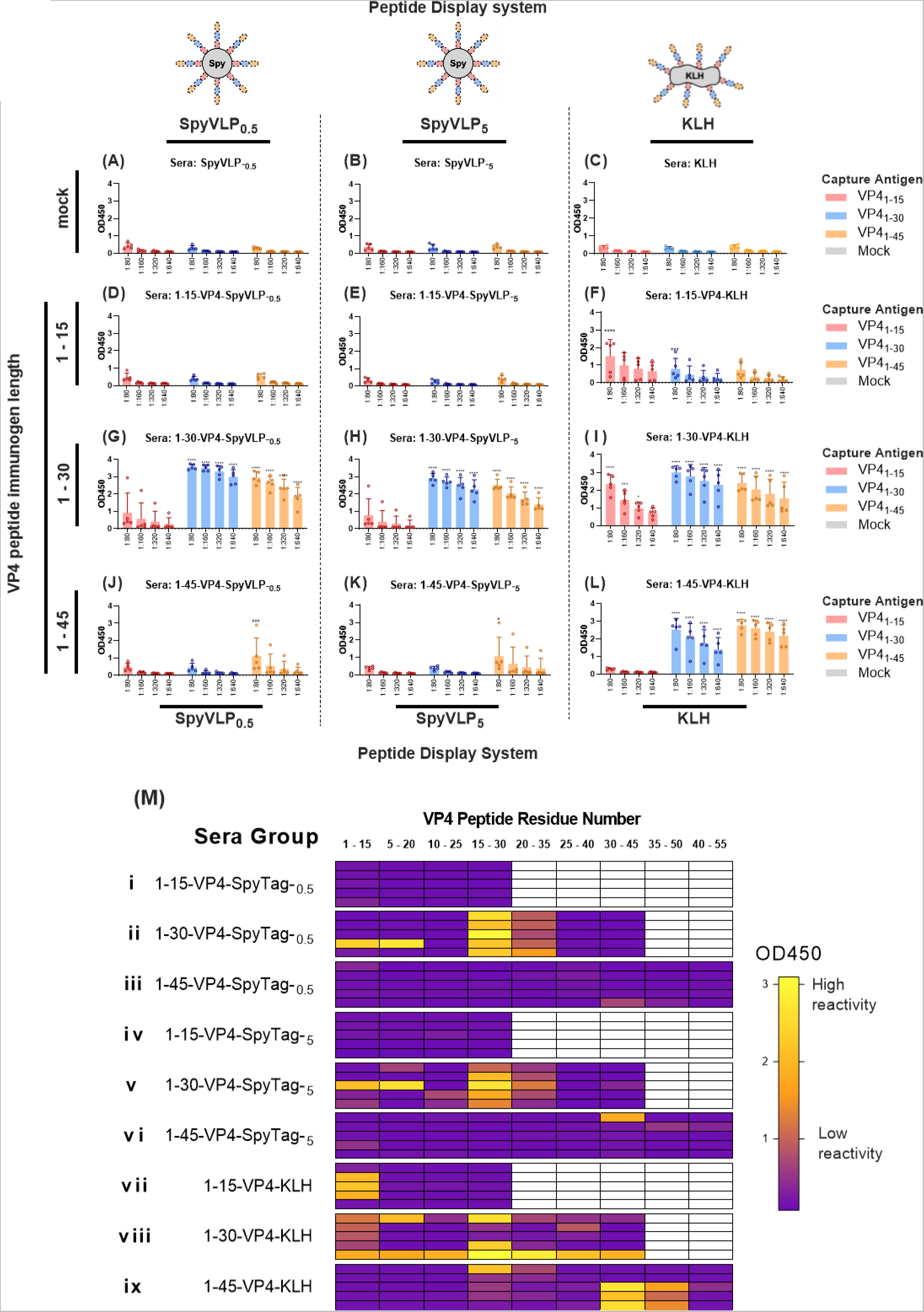
Reactvity of endpoint sera with VP4 capture peptides of different lengths. (A-L) ELISAs were used to determine the effect of peptide length on the reactivity with immune sera. Wells were coated with capture peptides corresponding to either 1-15-VP4 (red), 1-30-VP4 (blue) or 1-45-VP4 (yellow). ELISAs were performed for each sera group against each peptide using a dilution range of 1:80 to 1:640. Sera groups included SpyVLP-0.5 (A), SpyVLP-5 (B), KLH (C), 1-15-VP4-SpyVLP-0.5 (D), 1-15-VP4-SpyVLP-5 (E), 1-15-VP4-KLH (F), 1-30-VP4-SpyVLP-0.5 (D), 1-30-VP4-SpyVLP-5 (H), 1-30-VP4-KLH (I), 1-45-VP4-SpyVLP-0.5 (J), 1-45-SpyVLP-5 (K), 1-45-VP4-KLH (L). Absorbance was read at OD_450_. All ELISAs were carried out with anti-mouse IgG secondary antibody. Each point corresponds to a mouse (n=5 per vaccination group). Data are shown as mean ± SD. Statistical analysis was performed using one-way ANOVA and post-hoc analysis was performed using Dunnett’s test using the corresponding mock vaccination as the control mean. Significance threshold was set at p = 0.05. *p<0.05, ***p<0.0005, ****p<0.00005. (M) Heat map visualisation of epitope mapping. Each individual serum was analysed using ELISA with a scanning library of 15-mer peptides with 5 amino acid overlaps based on the first 55 N-terminal residues of RV16 VP4. Scanning peptides acted as the capture antigen. Amino acid residue numbers of the capture peptides are displayed above each column. ELISAs were performed for each sera group against each peptide using a dilution of 1:160. The sera group is labelled on the left: 1-15-VP4-SpyVLP-0.5 (i), 1-30 VP4-SpyVLP-0.5 (ii), 1-45-VP4-SpyVLP-0.5 (iii), 1-15-VP4-SpyVLP-5 (iv), 1-30-VP4-SpyVLP-5 (v), 1-45-VP4-SpyVLP-5 (vi), 1-15-VP4-KLH (vii), 1-30-VP4-KLH (viii), 1-45-VP4-KLH (ix). Absorbance was read at OD_450_. ELISAs were carried out with anti-mouse IgG secondary antibody. Each row corresponds to a single mouse (n=5 per vaccination group).

Analysis of the response against 1-15-VP4 peptide displayed by SpyVLP (1-15-VP4-SpyVLP) revealed that these sera did not react with any length of VP4 peptide, meaning that this immunogen did not induce an immune response that targets an epitope on free VP4 peptides (Figure 4 D,E,M i, iv). Given that the 1-15-VP4-SpyVLP-5 group was the most neutralising and the immunogen contained the 1-15-VP4 peptide, it is highly surprising that this serum has no detectable reactivity with free 1-15-VP4 peptide.

For 1-15-VP4-KLH, 3/5 of the sera did react with peptides 1-15-VP4, 1-30-VP4 and 1-45-VP4 peptides. However, the reactivity was highest with the 1-15-VP4 peptide (Figure 4F). Given that 1-30 and 1-45 peptides contain the equivalent sequence of the 1-15-VP4 peptide, these findings indicate that residues 1-15 may be presented in a different conformation in the longer peptides that reduces reactivity with the sera. This conformational change will likely either mask the epitope through steric hindrance, therefore making it less accessible to the sera, or will changes the conformation of the epitope so that the sera no longer recognise the peptide as efficiently. Further analysis by peptide scanning revealed that reactive 1-15-VP4-KLH sera only binds peptides that mapped to residues 1-15 (Figure 4 M vii). They did not react with peptides that mapped to residues 5-20 and above. This indicates binding to the first 5 amino acids of VP4 is important for the reactivity of these sera.

Analysis of the 1-30-VP4-SpyVLP sera revealed that 1/5 of the sera from both the 0.5 µg and 5 µg groups reacted with the 1-15-VP4 peptide. 5/5 in both groups reacted with 1-30 and 1-45 peptides, which indicates that the major epitope for 1-30-VP4-SpyVLP is located between residues 15-30 (Figure 4 G,H). Although reactivity was high with both the 1-30-VP4 and 1-45-VP4 peptide, reactivity was higher with the 1-30-VP4 peptide, indicating that this epitope may undergo a minor conformational change when the peptide length is increased from 30 to 45 amino acids (Figure 4 G,H). Further analysis by peptide scanning confirmed that the dominant epitope was located between residues 15-30 (Figure 4 M ii, v). Although the highest level of binding mapped to residues 15-30, these sera also mapped to residues 20-35, but the signal was lower. 2/5 of the sera from 1-30-VP4-SpyVLP-5 also showed reactivity with 10-25, but at very low levels; this reactivity was not observed in any 1-30-VP4-SpyVLP-0.5 sera (Figure 4 M ii, v). This region appears to form a minor epitope. Peptide scanning also revealed that sera from 1/5 mice from both the 1-30-VP4-SpyVLP-0.5 and 1-30-VP4-SpyVLP-5 groups were reactive with peptides 1-15 and 5-20, but not 10-25 (Figure 4 M ii, v). Signal was higher in the 5-20 peptide, indicating that this epitope is likely located between residues 5-15. Both sera that bound the 5-15 epitope also bound the dominant epitope at 15-30 described above (Figure 4 M ii, v). It is therefore not possible to distinguish between the signals for the two epitopes in the ELISAs with the longer peptides. Consequently we are unable to confirm if the 5-15 epitope is affected by increases in peptide length (Figure 4 G,H). However, since binding to the 5-15 epitope was only observed in 1/5 mice for each of the 1-30-VP4-SpyVLP groups, these results indicate that 5-15 is less dominant than the 15-30 epitope and it is therefore likely that length of the peptide does affect the exposure of this epitope.

Different results were observed in the 1-30-VP4-KLH sera. 1/5 of the sera were reactive with all peptides used in the peptide scan (Figure 4 M vii). This included peptide 30-45, which does not contain any VP4 residues present in the 1-30-VP4 immunogen. It does however contain the 6× lysines which are present in all peptides, this indicates that it does not bind a VP4 specific epitope and likely binds the lysines. This was the same serum that produced very high levels of 1-45-VP4 reactivity after the first immunisation (Figure 3I). Since this happened in only 1 out of the 30 mice immunised as part of this study, it means it a rare occurrence, but this reinforced the benefit of including the lysine-containing sequence in the peptide scanning analysis. The remaining 4 sera all reacted with peptides 1-15-VP4, 1-30-VP4 and 1-45-VP4, with peptide 1-30-VP4 showing the highest level of reactivity (Figure 4I). These data would be consistent with a major epitope located somewhere in residues 1-15, but where the conformation of this epitope is most dominant in the 1-30 peptide. This was supported by peptide scanning results: 4/4 sera reacted with peptides mapping to residues 1-15 and 1/4 bound 5-20 also (Figure 4 M viii). 2/4 sera could also bind peptides that mapped to residues 15-30 but, unlike the 1-30-VP4-SpyVLP, no strong reaction was observed with peptide 20-35. There was also weak reaction with peptides 25-40 in an individual serum. These data indicate that, unlike in 1-30-VP4-SpyVLP, for 1-30-VP4-KLH the dominant epitope may be within residues 1-15 and residues 15-30 only forms a minor epitope. These data indicate that the conformation of 1-30 is altered, depending on if 1-30 is conjugated to either SpyVLP or KLH.

Analysis of the 1-45-VP4-SpyVLP sera revealed that only 1/5 of the sera from both the 5 µg and 0.5 ug groups were strongly reactive with the 1-45-VP4 peptide and there was little reactivity in any sera with 1-15-VP4 or 1-30-VP4 peptides (Figure 4 J,K). This may indicate the presence of an epitope between residues 30-45. This was supported by peptide scanning, which revealed that these sera bound peptides that mapped to residues 30-45 (Figure 4 M iii, vi). No reactivity was observed with peptides 25-40 or 35-50, indicating that the epitope is located around residues 30-35 (Figure 4 M iii, vi). Also, the reactivity of the single positive 1-45-VP4-SpyVLP-0.5 serum was less reactive with the 30-45 peptide (Figure 4 M iii) than the 1-45 length peptide (Figure 4J), suggesting that the conformation of this epitope may not be optimal in the 30-45 peptide used in the peptide scanning. The peptide scan also revealed that individual sera had low levels of reactivity with peptides 1-15 and 35-50 (Figure 4M iii).

For 1-45-VP4-KLH sera, all sera were reactive with peptides 1-30-VP4 and 1-45-VP4, but the reactivity was strongest with the 1-45-VP4 peptide, while no reactivity was observed for the 1-15 peptide (Figure 4L). There are two scenarios consistent with this result: firstly that the sera could predominantly bind an epitope between residues 15-30, but the conformation is altered to become less dominant in the 1-30 peptide. Alternatively, the sera may bind multiple epitopes, one in 15-30 and another in 30-45. Peptide scans gave some clarification: despite 4/5 sera showing high reactivity with the 1-30 peptide (Figure 4L), only 1 had a strong signal with the 15-30 peptide (Figure 4M ix). This suggests that the sera that have high binding to the 1-30 peptide but lower levels of binding to the 15-30 peptide bind their target epitope in an area where the conformation is altered in the longer peptide. This conformation may be recapitulated poorly in the 15-mer peptides used in the scan. The peptide scan also showed that 3/5 of these sera reacted with peptides that mapped to 30-45 and 35-50, but not 25-40, indicating that these sera bind an epitope that spans residues 35-45 (Figure 4 M ix). 1/5 of the 1-45-VP4-KLH sera had no strong reactivity with any peptides in the scan, despite showing strong reactivity with 1-45 and weak reactivity with 1-30 (Figure 4 L,M ix). This peptide did have relatively weak levels of reactivity with peptide 15-30 (Figure 4M ix), indicating that the sera i binds the full-length peptide in a way that is dependent on the conformation of the full-length peptide and this conformation is recapitulated poorly in the 15-mer peptides used in the scanning.

Overall, these data indicate that the structure of VP4 is highly variable and the conformation of the epitopes that are displayed is highly dependent on the length of peptide and display system used. The antigenic conformation of these epitopes also appears to vary between free peptides, peptides conjugated to SpyVLP and peptides conjugated to KLH. Together these factors appear to influence what part of VP4 is targeted by the immune system. However, at present we were unable to identify where or what the neutralising sera from 1-15-VP4-SpyVLP immunised mice binds to. Given that these sera neutralise the virus they must bind the capsid. There are two potential explanations for these observations. We hypothesise that either the neutralising antibodies are not an IgG isotype which is the secondary antibody isotype we have used in all ELISA up to this point, or conjugation of the 1-15-VP4 peptide to the SpyVLP vector causes 1-15 to adopt a vastly different conformation to the free 1-15-VP4 peptide.

### Lack of peptide reactivity in 1-15-VP4-SpyVLP neutralising sera is not due to mis-matched isotype of detection antibody

All ELISAs up to this point used anti-IgG as the secondary antibody and we hypothesised that the lack of detectable reactivity between neutralising sera from 1-15-VP4-SpyVLP immunised mice and free VP4 peptide may be due to the neutralising sera containing non-IgG isotype antibodies. To determine the isotypes of VP4-specific antibodies in each sera, we screened the sera in ELISA using their cognate peptide as the capture, i.e. Peptide 1-15-VP4 was used as capture antigen for all sera raised against 1-15-VP4 peptides, peptide 1-30-VP4 was used as capture antigen for all sera raised against 1-30-VP4 peptides and peptides 1-45-VP4 was used as capture antigen for all sera raised against 1-45-VP4 peptides. We then screened the sera against a panel of different secondary antibodies, including anti-mouse IgA, IgM, IgG1, IgG2a, IgG2b or IG3 (Figure 5).

**Figure 5.**
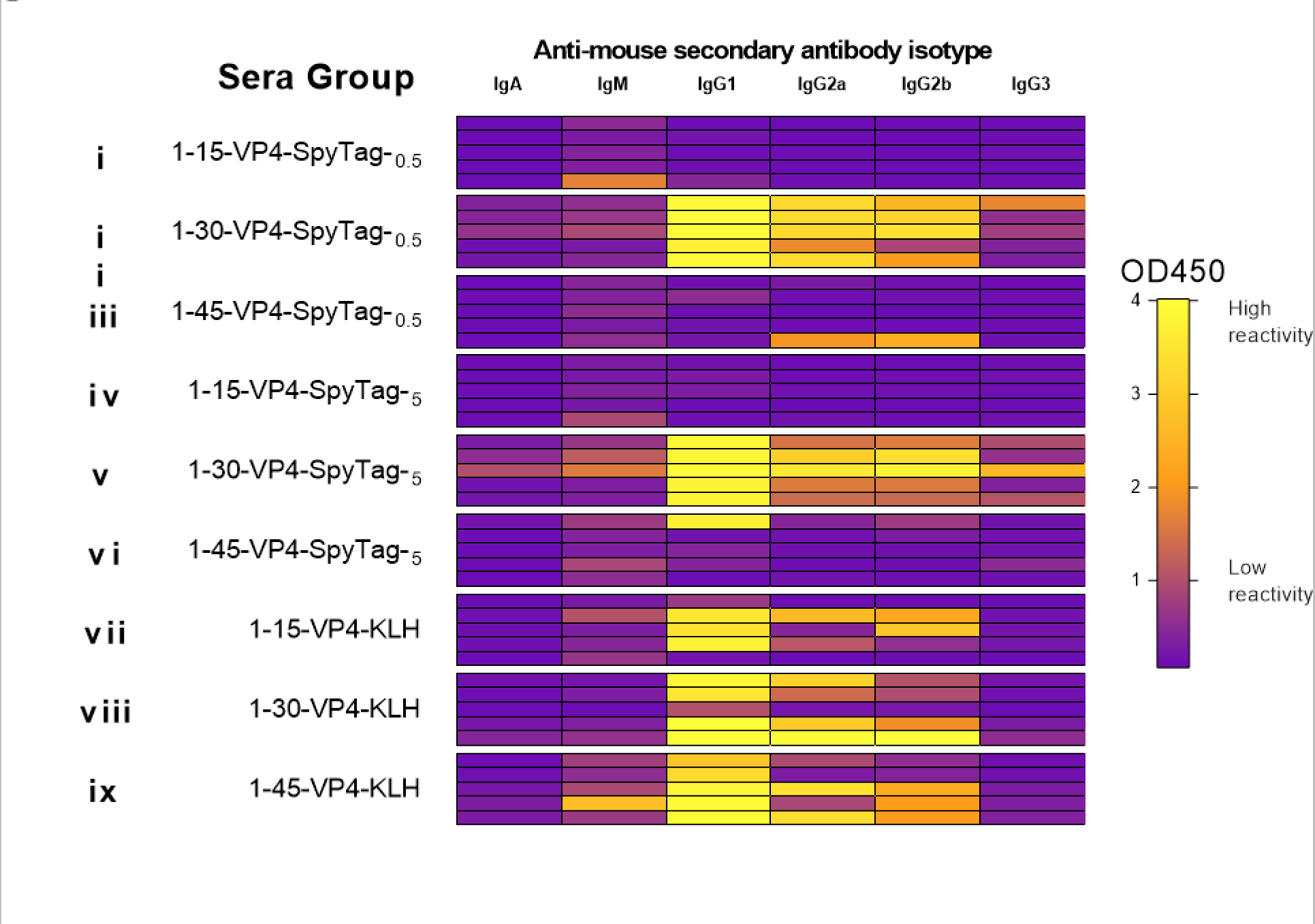
Heat map showing isotyping of sera groups. ELISAs were performed for each sera group with a different isotypes of mouse secondary antibodies (IgA, IgM, IgG1, IgG2a, IgG2b, IgG3). Data previously presented in figure 4 generated using a mixture of IgG isotypes (IgG mix) is included to act as a comparison. Peptide 1-15-VP4 was used as capture antigen for all sera raised against 1-15-VP4 peptides. Peptide 1-30-VP4 was used as capture antigen for all sera raised against 1-30-VP4 peptides. Peptide 1-45-VP4 was used as capture antigen for all sera raised against 1-45-VP4 peptides. ELISAs were performed for each sera group at a 1:160 dilution, sera group is labelled on the left, 1-15-VP4-SpyVLP-0.5 (i), 1-30 VP4-SpyVLP-0.5 (ii), 1-45 VP4-SpyVLP-0.5 (iii), 1-15-VP4-SpyVLP-5 (iv), 1-30 VP4-SpyVLP-5 (v), 1-45 VP4-SpyVLP-5 (vi), 1-15 VP4-KLH (vii), 1-30 VP4-KLH (viii), 1-45 VP4-KLH (ix). Absorbance was read at OD450. Each row corresponds to a single mouse (n=5 per vaccination group). Each column corresponds to a different secondary antibody isotype.

The results were consistent with existing data generated using the standard anti-IgG secondary (which is able to recognise IgG1, IgG2a, IgG2b and IgG3 antibodies). For example, sera from 1-15-VP4-SpyVLP-0.5 and 1-15-VP4-SpyVLP-5 immunised mice which were previously found not to be reactive with the standard anti-IgG secondary were also not reactive with secondary antibodies against any specific IgG subtype (Figure 5 i,iv). Antibodies previously shown to be reactive with a standard anti-IgG secondary were also highly reactive with at least one of the secondary antibodies against a specific IgG subtype. This includes sera from, 1-30-VP4-SpyVLP-0.5, 1-45-VP4-SpyVLP-0.5, 1-30-VP4-SpyVLP-5, 1-45-VP4-SpyVLP-5, 1-15-VP4-KLH, 1-30-VP4-KLH, 1-45-VP4-KLH immunised mice (Figure 5 ii, iii, v, vi, vii, viii, ix). Except for 1-45-VP4-SpyVLP-0.5 (Figure 5 iii), all these sera reacted most strongly with IgG1. This one sample that was previously showed reactivity with the standard IgG secondary serum had no detectable levels of reactivity with IgG1 but was reactive with IgG2a and IgG2b (Figure 5 iii).

Assessment of reactivity with an anti-mouse IgA secondary revealed that all sera had no or very little IgA (Figure 5).

Assessment of reactivity with an anti-mouse IgM secondary revealed that several sera such as 1-30-VP4-SpyVLP-5 and 1-45-VP4-KLH that had strong reactivity with anti-IgG secondaries also reacted with anti-IgM (Figure 5 v, ix). A single serum from 1-15-VP4-SpyVLP-0.5 reacted with anti-IgM, but this serum was not neutralising (Figure 5 i). However, none of the neutralising 1-15-VP4-SpyVLP-5 sera or the single neutralising serum from 1-15-VP4-SpyVLP-0.5 showed any substantial reactivity with anti-IgM (Figure 5 iv, i). Taken together, the lack of reactivity between the neutralising sera and VP4 peptides is not due to the neutralising sera belonging to a non-IgG isotype.

### Neutralising 1-15-VP4-SpyVLP sera only reacts with 1-15-VP4 peptide when conjugated to a display system

For serum to neutralise a RV-A16 infection, it must contain antibodies that bind the virus capsid. It is therefore surprising that neutralising sera from 1-15-VP4-SpyVLP immunised mice do not react with peptides that correspond to the VP4 capsid protein. We hypothesised that the neutralising 1-15-VP4-SpyVLP sera bind a native ‘virus-like’ conformational epitope within the first 15 N-terminal amino acids of VP4. This epitope may be induced by conjugation of the peptide to SpyVLP but would be poorly recapitulated in the free peptide.

We were unable to assess the ability of 1-15-VP4-SpyVLP-5 sera to bind the 1-15-VP4-SpyVLP immunogen due to high cross-reactivity with the SpyVLP presentation system. This caused high background reactivity and masked potential signal specific to 1-15-VP4. To overcome this issue, we tested if conjugation of VP4-1-15 peptide to an alternative peptide display system would reveal any peptide reactivity in the sera. For this we used an orthogonal peptide/protein pair that forms a spontaneous isopeptide bond, DogTag:DogCatcher (Keeble et al., 2022), to conjugate VP4-1-15 peptide to maltose binding protein (MBP). A subset of serum groups was selected (1-15-VP4-SpyVLP-5, 1-30-VP4-SpyVLP-5, 1-45-VP4-SpyVLP-5) to represent both neutralising and non-neutralising sera (Table 1). ELISA plates were coated with either conjugated 1-15-VP4-DogTag:DogCatcher-MBP or an unconjugated mix of DogCatcher-MBP and 1-15-VP4-SpyTag (peptide tagged with SpyTag; not compatible for conjugation to DogCatcher). Wells were then probed with sera from 1-15-VP4-SpyVLP-5, 1-30-VP4-SpyVLP-5 or 1-45-VP4-SpyVLP-5 immunised mice. As expected, the single 1-30-VP4-SpyVLP-5 sera that previously reacted with free 1-15 peptide (Figure 4H, 4M v) also reacted with the unconjugated 1-15-VP4 and DogCatcher-MBP (Figure 6A). For the remaining four 1-30-VP4-SpyVLP sera and 5/5 of the 1-15-VP4-SpyVLP-5 sera and 5/5 of the 1-45-VP4-SpyVLP-5 sera, reactivity with the unconjugated 1-15-VP4 and DogCatcher-MBP was at background levels (Figure 6). This demonstrates that these sera were unreactive with both free 1-15-VP4 peptide or DogCatcher-MBP. However, conjugation of 1-15-VP4-DogTag to DogCatcher-MBP enhanced reactivity for all sera that were non-reactive with the unconjugated mix of 1-15-VP4 and DogCatcher-MBP (Figure 6). This indicates that there are antibodies present in 1-15-VP4-SpyVLP-5, 1-30-VP4-SpyVLP-5 and 1-45-VP4-SpyVLP-5 sera, that are only reactive with the 1-15 peptide when it is conjugated to a display system and are not reactive with the free peptide. These sera likely react with a conformational epitope in 1-15-VP4 that only becomes accessible upon conjugation of the 1-15-VP4 peptide to SpyVLP or DogCatcher-MBP peptide presentation systems. However, given that this was observed in both neutralising and non-neutralising sera, this observation does not explain why some sera are neutralising while others are not.

**Figure 6.**
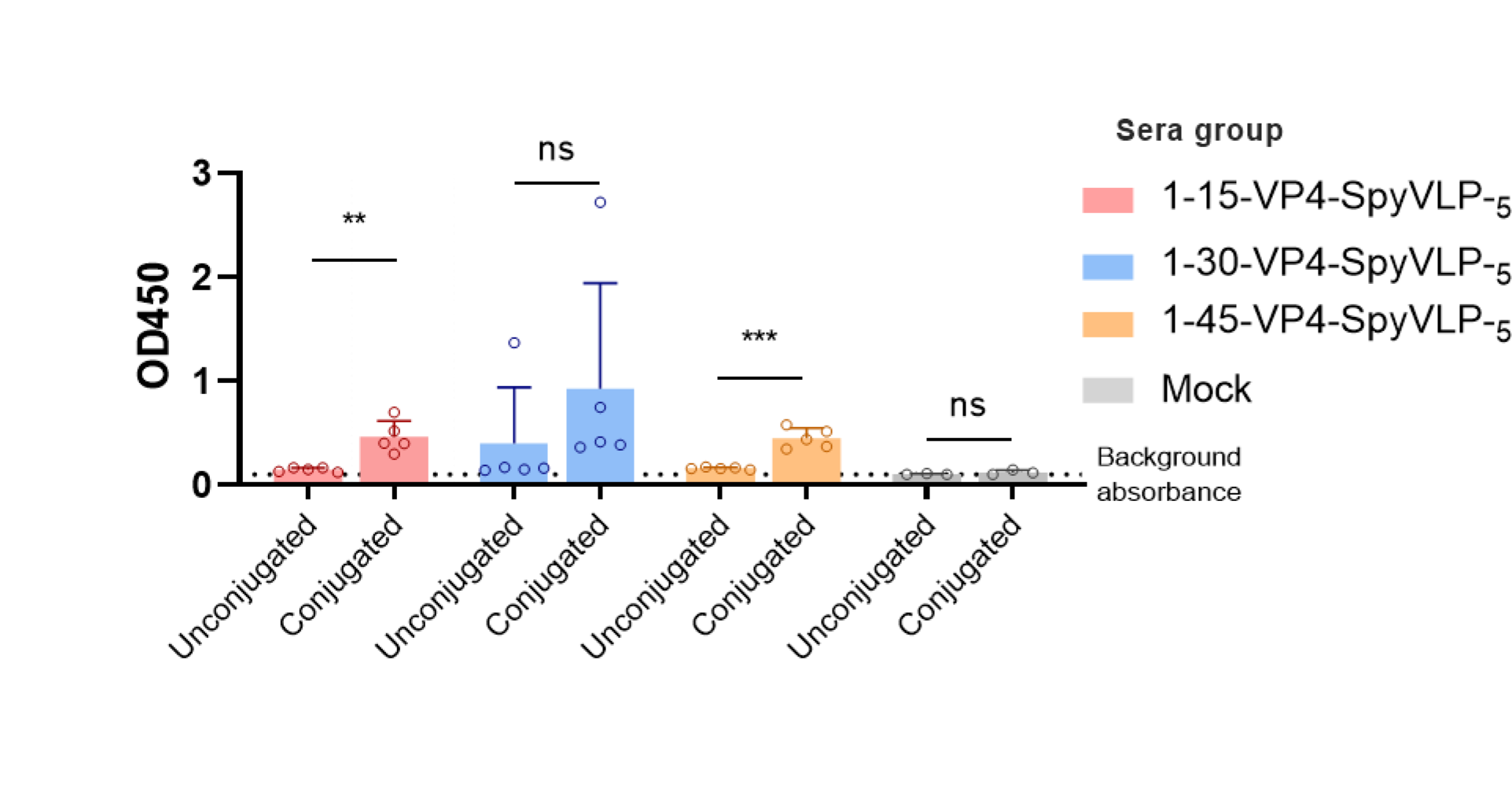
Reactivity of sera raised against SpyVLP-5 displayed peptides with either conjugated or unconjugated 1-15-VP4 peptide. ELISAs were performed to assess if conjugation of 1-15-VP4 peptide to DogCatcher-MBP affected its reactivity with sera from 1-15-VP4-SpyVLP-5, 1-30-VP4-SpyVLP-5 or 1-45-VP4-SpyVLP-5 immunized mice. Wells were coated with equimolar concentrations of either unconjugated or conjugated 1-15-VP4 solution to act as capture antigens. ELISAs were carried out using a 1:160 dilution with 1-15-VP4-SpyVLP-5 (red), 1-30-VP4-SpyVLP-5 (blue) or 1-45-VP4-SpyVLP-5 (yellow) or PBS as a negative control (grey). An IgG secondary antibody was used. Background absorbance is represented by a black dashed line. Absorbance was read at OD_450_. Each point corresponds to a mouse (n=5 per vaccination group). Data is shown as mean ± SD. Statistical analysis was performed using one-way ANOVA and post-hoc analysis was performed using Dunnett’s test using the corresponding mock vaccination as the control mean. Significance threshold was set at a = 0.05. *p<0.05, ***p<0.0005, ****p<0.00005.

### Neutralising 1-15-VP4-SpyVLP sera reacts with the RV-A16 capsid

Having established that most 1-15-VP4-SpyVLP-5, 1-30-VP4-SpyVLP-5 or 1-45-VP4-SpyVLP-5 sera only react with the 1-15-VP4 peptide when conjugated to a display system, we next sought to compare the ability of these sera to react with the RV-A16 capsid in ELISA using ICAM-1 (the RV-A16 receptor) to capture virus particles.

Assessment of reactivity between RV-A16 capsids and sera revealed that all sera tested showed OD readings above background. Mean readings for 1-15-VP4-SpyVLP-5 were 0.340 OD, average readings for 1-30-VP4-SpyVLP-5 were 0.260 OD and average readings for 1-45-VP4-SpyVLP-5 were 0.243 OD (Figure 7).

**Figure 7.**
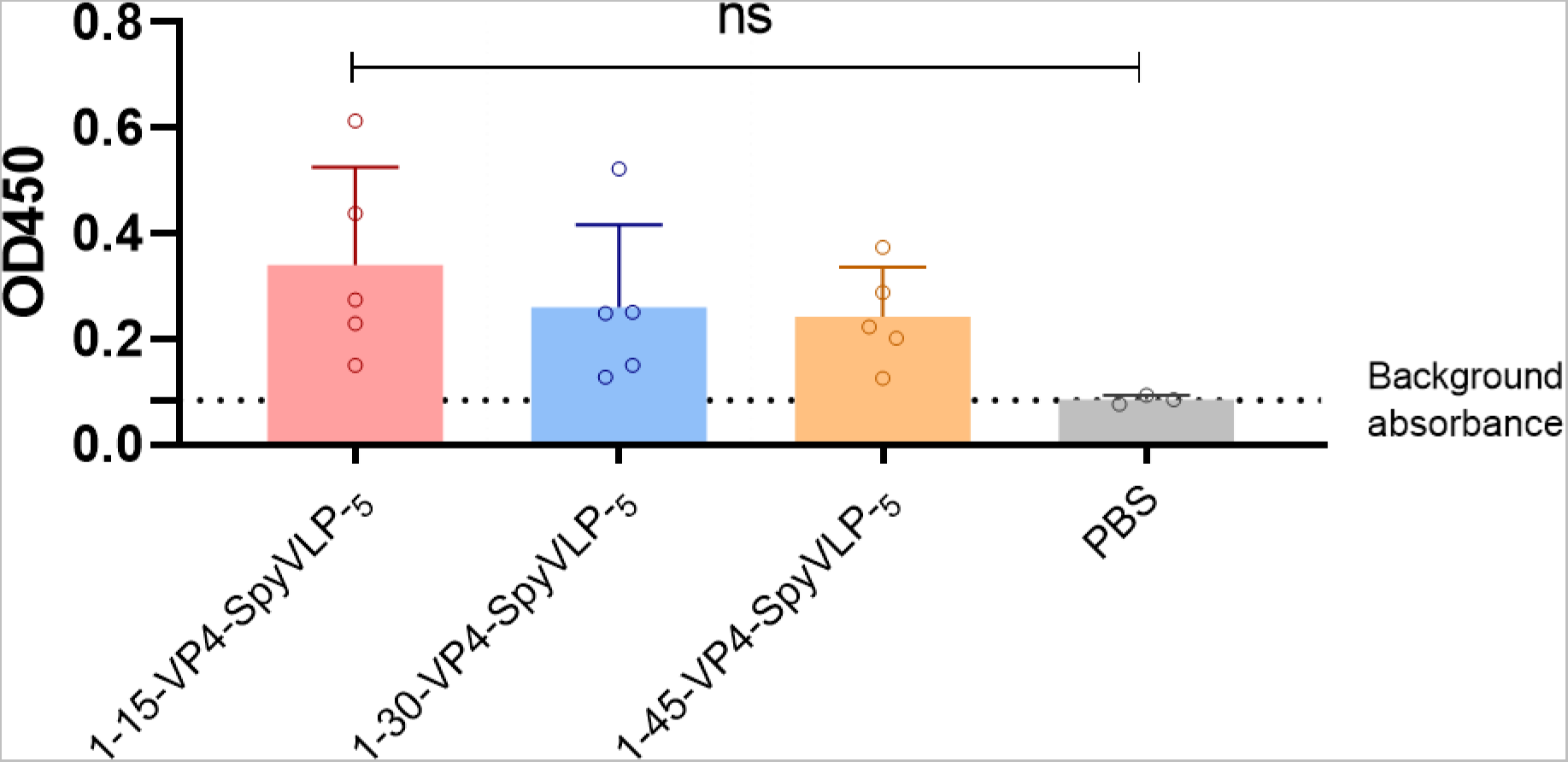
Reactivity of sera raised against SpyVLP-5 displayed peptides with RV-A16 capsids. ELISA was performed to assess ability of sera from 1-15-VP4-SpyVLP-5, 1-30-VP4-SpyVLP-5 or 1-45-VP4-SpyVLP-5 immunized mice to react with RV-A16 capsids. Plates were coated with ICAM-1 (the RV-A16 receptor) overnight. The following day RV-A16 antigen was captured on the plates using the receptors. ELISAs were carried out using a 1:160 dilution with 1-15-VP4-SpyVLP-5 (red), 1-30-VP4-SpyVLP-5 (blue) or 1-45-VP4-SpyVLP-5 (yellow), PBS negative control (grey). An IgG secondary antibody was used. Background absorbance is represented by a black dashed line. Absorbance was read at OD_450_. Each point corresponds to a mouse (n=5 per vaccination group). Data are shown as mean ± SD. Statistical analysis was performed using one-way ANOVA and post-hoc analysis was performed using Dunnett’s test using the corresponding mock vaccination as the control mean. Significance threshold was set at a = 0.05. *p<0.05, ***p<0.0005, ****p<0.00005.

This demonstrates that immunisation with 1-15-VP4-SpyVLP-5, 1-30-VP4-SpyVLP-5 or 1-45-VP4-SpyVLP-5 can generate antibodies that can react the RV-A16 capsid. Of these immunogens, 1-15-VP4-SpyVLP produced sera that was most reactive with the RV-A16 capsid. This correlated with only these sera having neutralising activity (Figure 2B). However, higher capsid reactivity does not entirely explain neutralisation: although 1-15-VP4-SpyVLP-5 had the highest average reactivity of the three groups, individual non-neutralising sera from groups 1-30-VP4-SpyVLP-5 and 1-45-VP4-SpyVLP-5 had higher reactivity with the capsid than individual neutralising sera from the 1-15-VP4-SpyVLP-5 group (Figure 7). This indicates there is a more nuanced mechanism of neutralization than simply binding the capsid.

## Discussion

In the current study we sought to identify conditions to create a RV VP4 immunogen that generates the greatest possible neutralising response. We compared three different lengths of VP4 peptide (1-15-VP4, 1-30-VP4 and 1-45-VP4) conjugated to two different display system (SpyVLP and KLH). We showed that the combination of the 1-15-VP4 peptide and SpyVLP display system (1-15-VP4-SpyVLP) produced the highest levels of neutralising antibodies. Other immunogens either produced a response that was 8 times lower than 1-15-VP4-SpyVLP or no neutralising response at all. The 1-15-VP4-SpyVLP immunogen also produced the most consistent response: a 5 µg dose of 1-15-VP4-SpyVLP produced a neutralising response in all 5 mice, while no other immunogen achieved this.

Mapping reactivity of the resulting sera to free peptides identified several antigenic sites in RV-A16 VP4, including sites that map to residues 1-15 and 15-30. However, antibody binding to these sites did not correlate with neutralisation. This was surprising given that previous studies with other picornaviruses identified neutralising antibodies that reacted with VP4 residues 7-15 in RV-B14 and residues 5-12 and 25-35 in EV-A71 (Katpally et al., 2009; Phanthong et al., 2020; Zhao et al., 2013). Instead, in our study VP4-specific antibody neutralization of RV-A16 was dependent on binding a virus-like conformational epitope in 1-15-VP4. We suspect that 1-15-VP4-SpyVLP produced the highest level of neutralising antibodies because it most accurately recapitulates a virus-like conformation of the 1-15 epitope. It is possible that antibodies that target other sites may be neutralising, but they did not achieve a high enough titre in this study.

Further detailed mapping of the different sera produced in this study revealed that the antigenic conformation of VP4 is highly variable and is influenced by peptide length and display system. Examples of peptide length influencing antigenic conformation is demonstrated in the difference reactivities of some of the sera with different lengths of peptide. The 1-15-VP4-KLH sera reacted with free 1-15-VP4 peptide but did not react with equimolar concentrations of 1-30 and 1-45 peptides (Figure 4F). This indicates that residues 1-15 may be presented in a different conformation in the longer peptides, that reduces reactivity with the 1-15-VP4-KLH sera. Furthermore, 1-30-VP4-SpyVLP generate a strong response directed against residues 15-30 of VP4 (Figure 4 G, H, M ii, v). However, sera generated in response to the 1-45-VP4-SpyVLP, which also contains the 15-30 sequence, had no detectable response against the peptide mapping to this region (Figure 4 J, K, M vi). This indicates that presentation of the 15-30 region of VP4 differs between 1-45-VP4-SpyVLP and 1-30-VP4-SpyVLP.

By comparing reactivities of the 1-30-VP4-SpyVLP and 1-30-VP4-KLH sera, we demonstrate that display system also influences the antigenic conformation. Most sera from mice immunised with peptides presented on KLH (1-15-VP4-KLH or 1-30-VP4-KLH) were reactive with free 1-15-VP4 peptide. However, sera from mice immunised with peptides presented on SpyVLP (1-15-VP4-SpyVLP or 1-30-VP4-SpyVLP) were unreactive with free 1-15-VP4 peptide, but they were reactive with the peptide after conjugation to a DogCatcher-MBP display system. These data indicate that the 1-15-VP4 peptide undergoes a conformation change upon conjugation to the SpyVLP or DogCatcher-MBP but not KLH. The difference we detected here however do not explain the cause of the superior neutralisation ability of 1-15-VP4-SpyVLP sera, since non-neutralising sera from 1-30-VP4-SpyVLP and 1-45-VP4-SpyVLP produced similar levels of reactivity with the conjugated 1-15-VP4 antigen. We therefore suspect that there are further differences in conformation between 1-15-VP4-SpyVLP and 1-30-VP4-SpyVLP/1-45-VP4-SpyVLP which we cannot detect in our ELISA.

Previous studies with other picornaviruses also indicated that peptide length and display system influence antigenic conformation of VP4. Immunisations with RV-B14 peptides corresponding to VP4 1-24 generated neutralising antibodies that were reactive with free VP4 1-24 peptides, and 15-mer peptides that contained residues 7-15 of VP4. In contrast, immunisation with peptides corresponding to residues 1-30 generated neutralising antibodies that were reactive with full length VP4, but which had no detectable reactivity with the 1-24-VP4 peptide or any 15mer peptides containing residues 7-15 (Katpally et al., 2009). This demonstrates an increase in peptide length of just 6 amino acids is enough to completely change the antigenicity of VP4.

For EV-A71, mice were immunised with a Hepatitis B Virus core fusion protein VLP containing the N-terminal 20 amino acids of EV-A71 VP4 and this generated neutralising antibodies that bound the 5-12 region of VP4 (Zhao et al., 2013). However, in a separate study, a phage display screen using full length VP4 as bait only generated antibodies that bound between residues 25-34 (Phanthong et al., 2020; Zhao et al., 2013). The phage display screen did not generate antibodies that bound the 5-12 epitope, which indicates that the 5-12 epitope is not available in the full-length protein. These data highlight that the antigenic conformation of VP4 is highly variable, which has important implications on the design of a future VP4-based vaccine. The ability of VP4 to adopt multiple conformations is likely related to the multiple biological functions required of this protein in capsid assembly, stability and uncoating. Structural studies of antibody-antigen complexes would facilitate a better understanding of the conformation of peptides and determine optimum conformations for neutralisation.

While the current study has focussed on neutralising antibody responses, it is possible that the immunogens tested in this study may produce protective T-cell responses or non-neutralising antibodies may initiate other modes of protection, such as antibody-dependent cellular phagocytosis (ADCP). Previous studies have shown that antibodies targeting the C-terminus of RV VP1, which like VP4 is an internal epitope, are non-neutralising but can protect via ADCP (Behzadi et al., 2020). Furthermore, antibodies targeting the C-terminus of VP4 may contribute to a protective T-cell response in mice (Glanville et al., 2013; Narean et al., 2019).

In conclusion, this study has demonstrated that presentation of the 1-15 region of VP4 in a specific antigenic conformation is important for the generation of neutralising VP4 antibodies. However, the antigenic conformation of VP4 varies greatly between peptides of different lengths and identical peptides presented on different display systems. Furthermore, even the best immunogen created in this study produced a relatively low level of neutralising antibodies. This highlights the complexities of creating an effective response against VP4. Future efforts to develop a VP4 vaccine would need to focus on improving neutralising antibody titres and investigate the potential for non-neutralising responses to VP4 to provide protection against RV infection. This work could also impact the design of other peptide vaccines, since a highly variable antigenic conformation may be a common characteristic of peptides and not unique to VP4 peptides.

## Acknowledgements

We thank the Pirbright Institute animal services team for carrying out the mouse immunisations.

## Materials and methods

### Peptides

RV-A16 VP4 peptides (Table 2) were synthesised by Peptide Synthetics. Peptides were dissolved at 33.3 mM in dimethyl sulfoxide (D2650, Sigma). Stocks were then aliquoted and stored at -20 °C until further use. Peptides were either conjugated to display vectors to create immunogens or used as capture antigens in ELISAs. Peptides used as capture antigens for ELISA were diluted in 0.05 M carbonate-bicarbonate buffer to a concentration of 2.4 mM.

**Table 2.**
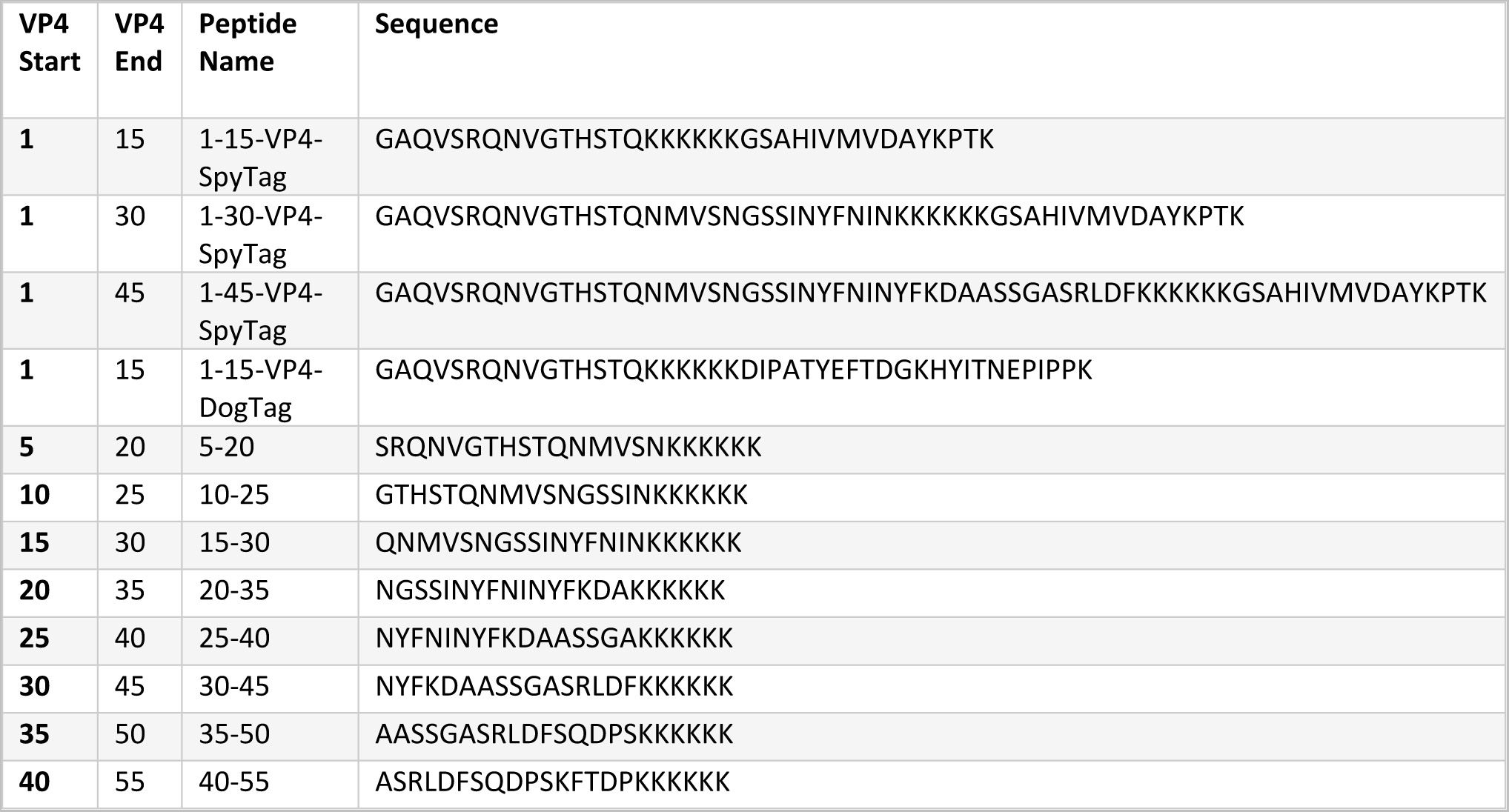
Peptides used in the study.

### Expression and purification of SpyCatcher003-mi3 and DogCatcher-MBP display systems

pET28a-SpyCatcher003-mi3 (GenBank MT945417, Addgene 159995) (Rahikainen et al., 2021) and pET28a-AviTag-DogCatcher-MBP (GenBank MZ365293, Addgene 171928) (Keeble et al., 2022) were transformed into *E. coli* BL21(DE3)-RIPL cells. They were grown on LB-Agar plates with 50 μg/mL kanamycin for 16 h at 37 °C. A single colony was used to inoculate 10 mL LB medium containing 50 μg/mL kanamycin and grown for 16 h at 37 °C with shaking at 200 rpm. This culture as added to 1 L LB + 0.8% (w/v) glucose containing 50 μg/mL kanamycin and incubated at 37 °C and 200 rpm shaking until OD_600_ reached 0.6. Cultures were induced with 0.5 mM isopropyl β-D-1-thiogalactopyranoside (IPTG). For SpyCatcher003-mi3, cells were grown at 22 °C with shaking at 200 rpm for 16 h. For AviTag-DogCatcher-MBP, cells were grown at 30 °C with shaking at 200 rpm for 4 h. Cultures were pelleted by centrifugation at 4,000 g.

AviTag-DogCatcher-MBP pellets were resuspended in 20 mL Ni-NTA buffer (50 mM Tris-HCl pH 7.8 containing 300 mM NaCl) and SpyCatcher003-mi3 pellets were resuspended in 20 mL 20 mM Tris-HCl, 300 mM NaCl, pH 7.9 at 25 °C. In both cases the pellets were supplemented with 0.1 mg/mL lysozyme, 1 mg/mL complete mini EDTA-free protease inhibitor (Roche), and 1 mM phenylmethanesulfonyl fluoride (PMSF) and incubated for 30 min. An Ultrasonic Processor equipped with a microtip (Cole-Parmer) was used to perform sonication on ice (4 times for 60 s, 50% duty-cycle). Cell debris were cleared by centrifugation at 35,000 g for 45 min at 4 °C.

AviTag-DogCatcher-MBP was purified by nickel-nitrilotriacetic acid (Ni-NTA) affinity chromatography. Ni-NTA agarose (Qiagen) was packed in an Econo-Pac Chromatography Column (Bio-Rad). It was washed with 2 × 10 CV of Ni-NTA buffer. The bacterial supernatant was incubated in the Ni-NTA column for 1 h at 4 °C with rolling. The supernatant was allowed to flow through by gravity, before being washed with 2 × 10 CV of Ni-NTA wash buffer (10 mM imidazole in Ni-NTA buffer). The resin was incubated with Ni-NTA elution buffer (200 mM imidazole in Ni-NTA buffer) for 5 min before eluting by gravity. A total of six 1 CV elutions were performed. Elution fractions were assessed by SDS-PAGE with Coomassie staining, pooled, and dialyzed for 16 h against 1,000-fold excess PBS before being stored at -80 °C.

SpyCatcher003-mi3 was purified by ammonium sulfate precipitation. 170 mg of ammonium sulfate was added per mL of cleared lysate. The mixture was incubated at 4 °C for 1 h with stirring at 100 rpm to precipitate the particles. The solution was centrifuged for 30 min at 30,000 g at 4 °C and the pellet was resuspended in 10 mL mi3 buffer (25 mM Tris–HCl, 150 mM NaCl, pH 8.0) at 4 °C. The resuspended pellet was filtered sequentially through 0.45 µm and 0.22 µm syringe filters (Starlab).

The filtrate was dialyzed for 16 h against 1,000-fold excess mi3 buffer using a 100 kDa molecular weight cutoff Spectra/Por-3 dialysis membrane (Spectrum Labs). The dialysed particles were centrifuged at 17,000 g for 30 min at 4 °C and filtered through a 0.22 µm syringe filter. The filtrate was loaded onto a HiPrep Sephacryl S-400 HR 16-600 column (GE Healthcare), which had been equilibrated with filtered mi3 buffer using an ÄKTA Pure 25 system (GE Healthcare). The protein was run over the column at 0.1 mL/min while collecting 1 mL elution factions. The fractions containing the purified SpyCatcher003-mi3 were pooled and concentrated using a Vivaspin 20 100 kDa molecular weight cut-off centrifugal concentrator (GE Healthcare).

SpyCatcher003-mi3 was endotoxin-depleted using Triton X-114 phase separation. For this procedure, all microcentrifuge tubes were pyrogen-free and all pipette tips were filtered. 1% (v/v) Triton X-114 was added to SpyCatcher003-mi3 samples and the solution was mixed by gentle pipetting. The samples were incubated on ice for 5 min. The solution was transferred to a ThermoMixer (Eppendorf) at 37 °C and incubated for 5 min and centrifuged for 1 min at 16,000 g at 37 °C. The top phase was transfered into a fresh tube. This procedure was repeated for a total of three times. After the third round, an additional incubation at 37 °C for 5 min, followed by centrifugation at 16,900 g for 2 min at 37 °C, was performed to remove residual Triton X-114. The endotoxin content of all recombinant antigens was measured using the Limulus amebocyte lysate (LAL)-based Chromogenic Endotoxin Quantitation Kit (Thermo Fisher) following manufacturer’s instructions. All particles were below the acceptable levels of 20 EU/mL (Brito & Singh, 2011). The concentration of endotoxin-depleted particles was measured using bicinchoninic acid (BCA) assay (Pierce) and then stored at -80 °C.

### Conjugation of VP4 peptides to display systems

VP4 peptides 1-15-VP4-SpyTag, 1-30-VP4-SpyTag or 1-45-VP4-SpyTag at 10 µM were conjugated to SpyVLP at 10 µM in PBS at 25 °C for 18 h. Possible aggregates were then removed by centrifugation at 16,900 g for 30 min at 4 °C.

1-15-VP4-DogTag at 10 mM was conjugated to DogMBP at 10 mM in PBS at 25 °C for 2 h.

Conjugation of KLH to myristoylated 1-15-VP4, 1-30-VP4, and 1-45-VP4 peptides was performed by Peptide Synthetics.

### SDS-PAGE

Samples were mixed with reducing 6× loading dye (0.23 M Tris–HCl, pH 6.8, 24% (v/v) glycerol, 120 μM bromophenol blue, 0.23 M SDS, 0.2 M dithiothreitol) and resolved on 12% SDS–PAGE. Gels were then stained with InstantBlue Coomassie (Expedion) and imaged using a ChemiDoc XRS imager (Bio-Rad).

### DLS

Samples were centrifuged for 30 min at 16,900 × g at 4 °C to pellet possible aggregates. Before each measurement, the quartz cuvette was incubated in the instrument for 5 min to stabilise the sample temperature. Samples were measured at 100 μg/mL SpyVLP. 30 μL of sample was measured at 20 °C using an Omnisizer (Victotek) with 20 scans of 10s each. The settings were 50% laser intensity, 15% maximum baseline drift, and 20% spike tolerance.

### Cells and virus

H1 HeLa cells (CRL-1958, ATCC) were propagated in 10% (v/v) foetal bovine serum (FBS) (26140087, Gibco) + 1% (v/v) penicillin-streptomycin (P/S) (15140122, Gibco) in Dulbecco’s Modified Eagle Medium (DMEM) (41965039, Gibco) at 37 °C and 5% (v/v) CO_2_. 80-90% confluent cells were split by washing with sterile PBS and incubation with 0.25% (v/v) trypsin-EDTA (25200056, Gibco) for 3 min. Trypsin was inactivated with 10% (v/v) FBS-DMEM and cells were pelleted at 900 x g for 5 min at 22°C. Supernatants were discarded and pellets were resuspended and diluted in 10% (v/v) DMEM for use in subsequent experiments. RV-A16 (VR-283, ATCC) were propagated by infection of 80-90% confluent H1 HeLa cells. Cell lysates were collected upon death of 90% of the monolayer.

### Virus purification

Cell lysates were collected upon death of 90% of the monolayer and cell debris was pelleted by centrifugation at 900 x g for 5 min at 25 °C and resuspended in 1% (v/v) NP-40. Cell debris was then freeze-thawed three times at -20 °C and then pelleted by centrifugation at 3000 x g for 10 min at 22°C. Supernatants were pooled and precipitated by incubation in 50% saturated ammonium sulfate perception at 4 °C overnight, followed by centrifugation at 4,000 x g for 1 h at 4 °C. Supernatant was discarded and the precipitate was resuspended in PBS with 1% (v/v) NP-40. Precipitate was pelleted though a 30% (w/v) sucrose cushion at 120,000 × *g* (using a Beckman SW32 Ti rotor) for 3.5 h at 4°C. The pellet was resuspended in PBS and clarified by differential centrifugation. The supernatant was purified through a 15 to 45% (w/v) sucrose density gradient by ultracentrifugation at 160,000 × *g* (using a Beckman SW40 Ti rotor) for 2.5 h at 4 °C. Gradients were fractionated and peak fractions corresponding to virions were identified by OD at 260/280 nm.

### Mouse immunisations and sampling

Mouse experiments were performed according to the UK Animals (Scientific Procedures) Act, Project Licence (PP8335202) and approved by The Pirbright Institute Animal Request Review Body. All experiments were done in compliance with the ethical regulations for animal testing and research. All conditions were in accordance with or surpassed the UK Home Office ethical and welfare guidelines. Female BALB/c mice (∼9 weeks old at the time of first immunisation) were obtained from Envigo (now Inotiv). Mice were housed in large mouse breeding cage rather than a M1 mouse cage and bedding was corn cob. Mice were provided with mouse swings and red plastic items to climb on/hide in as environmental enrichment (e.g. igloos and tubes). Mice were handled using clear tubes rather than tail handling. Mice were fed on RM1 (e) Maintenance diet with water ad libitum. The mouse room was kept at 20.0 – 21.4 °C with 57.7%–61.6% humidity. The light cycle was 12 h on, 12 h off with a lux reading of at cage level of ∼ 25 Lux.

To prepare the SpyVLPs for vaccination, 10 μM SpyTag-VP4 was conjugated with 10 μM of SpyCatcher003-mi3 in PBS at 25 °C for 18 h. The reaction was centrifuged for 30 min at 17,000 × g at 4 °C to remove potential aggregates. Doses were diluted to either 10 µg/ml (SpyVLP-VP4-0.5) or 100 µg/ml (SpyVLP-VP4-5) in sterile PBS. For Spy-VLP immunisations, fresh SpyCatcher-conjugated immunogens were prepared for each boost. For KLH immunisations, unfrozen immunogen was used for the first immunisation and for the boosts aliquots of immunogen that had been subject to one freeze-thaw were used. KLH doses were diluted to 1,000 µg/ml in sterile PBS. Before subcutaneous (SC) immunisation, each sample was mixed 1:1 with MagicMouse (Gentaur or Invivogen) and immunised with 100 µL this solution. This created a final dosage of either 0.5 µg for SpyVLP-0.5, 5 µg for SpyVLP-VP4-5 or 50 µg for KLH.

Mice were prime-boost-boost vaccinated subcutaneously with either 1-15-VP4-SpyVLP-0.5, 1-30-VP4-SpyVLP-0.5, 1-45-VP4-SpyVLP-0.5, SpyVLP-0.5, 1-15-VP4-SpyVLP-5, 1-30-VP4-SpyVLP-5, 1-45-VP4-SpyVLP-5, SpyVLP-5, 1-15-VP4-KLH, 1-30 VP4-KLH, 1-45 VP4-KLH or KLH. Each vaccination group consists of 5 mice. Bleeds were taken from the tail 10 days and 24 days post immunisation. At day 42, mice were anaesthetised and terminated by cardiac puncture terminal bleeding. The anaesthetic used was a cocktail of 300 mg/kg Ketamine and 3mg/kg Medetomidine dosage, according to mouse weight. To prepare the anaesthetic cocktail, equal volumes of 100 mg/mL Ketamine and 1mg/mL Medetomidine were mixed thoroughly. Volume administered to each mouse was based on their weight, e.g. for a 25 g mouse, 0.15 mL of this cocktail was given. The ketamine/medetomidine anaesthetic ensured mice did not feel the cardiac puncture and that they have a high heart rate. This enables a higher volume of blood to be harvested from each mouse. Mice were confirmed to be unresponsive prior to cardiac puncture. Bloods were collected into 1.5 mL microcentrifuge tubes and allowed to clot at 22 °C for 1 h, before spinning down at 10,000 × g for 5 min at 22 °C. The clarified sera were stored at -20 °C.

### ELISA

96-well plates were coated overnight with 50 μL capture antigen (peptides 2.4mM, peptide/display vector 2.4 mM/2.4 mM, ICAM-1 (supplied by PEPRO TECH) 0.2 µg/mL) diluted in 0.05 M carbonate-bicarbonate buffer overnight at 4 °C. Wells were washed 5 times with PBS with 0.1% (v/v) Tween (PBS-T) and blocked with 5% blocking buffer (5% (w/v) skimmed milk (Marvel) + PBS-T) for 1 h at 22 °C. For ICAM-1-containing wells, an additional incubation was conducted. Here wells were incubated with 100 µL of 0.07 µg/mL purified RV-A16 in 1% (w/v) bovine serum albumin for 1 h to capture the virus and then washed 5 times with PBS-T at 22 °C. All capture antigen wells were incubated with sera or controls diluted in 1% skimmed milk in PBS-T at either a 1:160 dilution or twofold serial dilutions spanning 1:80 to 1:640 for 1 h at 22 °C. Wells were washed with PBS-T and incubated with HRP-conjugated antibodies for 1 h at 22 °C. Wells were washed with PBS-T and incubated with 100 μL 1-Step Ultra TMB-ELISA (34029, Thermo) for 8 min and stopped with 0.16 M H_2_SO_4_. Colorimetric change was quantified using a GloMax microplate reader at OD_450_.

### Virus neutralisation test (VNT)

Serially diluted sera were incubated with 1.4 × 10^4^ PFU/mL RV-A16 in 4% (v/v) FBS + 1% (v/v) penicillin/streptomycin DMEM at 4°C overnight. Sera + RV16 mixtures were then added to 96-well plates containing 80-90% confluent H1-HeLa cells in 4% (v/v) FBS + 1% (v/v) penicillin/streptomycin in DMEM. Plates were incubated in 37 °C with 5% (v/v) CO_2_ for 3 days. Wells were then stained with 100 μL crystal violet fixing and staining solution (2.5 g crystal violet + 50 mL ethanol + 380.6 mL PBS + 70mL 36% (w/v) paraformaldehyde) for 10 min 22 °C. Plates were washed with water and allowed to air dry at 22 °C. Wells were incubated with 100 μL 1% (w/v) sodium dodecyl sulfate (SDS) in dH_2_O for 15 min 22 °C in an orbital shaker. Resuspended crystal violet was quantified using a GloMax microplate reader at OD_560_.

### Statistical analysis

Statistical analysis and data visualisation were performed using a licensed GraphPad Prism (version 9.1.2). OD_450_ and OD_560_ obtained were subtracted from blank control wells and plotted as such. Statistical analysis was performed using one-way ANOVA for all experiments with the significance threshold set at α = 0.05. Post-hoc analysis was performed based on the type of comparisons performed. For data compared to a single control setup, we performed Dunnett’s test. For data where the change in signal per mouse is of interest, we performed paired t-tests. For data where multiple comparisons are performed, we utilised two-way ANOVA and post-hoc analysis was performed using Sidak’s multiple comparison test. Data were presented as mean ± 1 S.D. when applicable.

